# Rif1 inhibits replication fork progression and controls DNA copy number in Drosophila

**DOI:** 10.1101/346650

**Authors:** Alexander Munden, Zhan Rong, Rama Gangula, Simon Mallal, Jared T. Nordman

## Abstract

Control of DNA copy number is essential to maintain genome stability and ensure proper cell and tissue function. In Drosophila polyploid cells, the SNF2-domain-containing SUUR protein inhibits replication fork progression within specific regions of the genome to promote DNA underreplication. While dissecting the function of SUUR’s SNF2 domain, we identified a physical interaction between SUUR and Rif1. Rif1 has many roles in DNA metabolism and regulates the replication timing program. We demonstrate that repression of DNA replication is dependent on Rif1. Rif1 localizes to active replication forks in an SUUR-dependent manner and directly regulates replication fork progression. Importantly, SUUR associates with replication forks in the absence of Rif1, indicating that Rif1 acts downstream of SUUR to inhibit fork progression. Our findings uncover an unrecognized function of the Rif1 protein as a regulator of replication fork progression.

## INTRODUCTION

Accurate duplication of a cell’s genetic information is essential to maintain genome stability. Proper regulation of DNA replication is necessary to prevent mutations and other chromosome aberrations that are associated with cancer and developmental abnormalities (Jackson et al., 2014). DNA replication begins at thousands of cis-acting sites termed origins of replication. The Origin Recognition Complex (ORC) binds to replication origins where, together with Cdt1 and Cdc6, it loads an inactive form of the MCM2-7 replicative helicase (Bell and Labib, 2016). Inactive helicases are phosphorylated by two key kinases, S-CDK and Dbf4-dependent kinase (DDK), which results in the activation of the helicase and recruitment of additional factors to form a pair of bi-directional replication forks emanating outward from the origin of replication (Siddiqui et al., 2013). Although many layers of regulation control the initiation of DNA replication, much less in known about how replication fork progression is regulated.

In metazoans, replication origins are not sequence specific and are likely specified by a combination of epigenetic and structural features (Aggarwal and Calvi, 2004; Cayrou et al., 2011; Eaton et al., 2011; Mesner et al., 2011; Miotto et al., 2016; Remus et al., 2004). Furthermore, replication origins are not uniformly distributed throughout the genome. The result of non-uniform origin distribution is that, in origin-poor regions of the genome, a single replication fork must travel great distances to complete replication. If a replication fork encounters an impediment within a large origin-less region of the genome, then replication will be incomplete, resulting in genome instability (Newman et al., 2013). In fact, origin poor regions of the genome are known to be associated with chromosome fragility and genome instability (Debatisse et al., 2012; Durkin and Glover, 2007; Letessier et al., 2011; Norio et al., 2005). This highlights the need to regulate both the initiation and elongation phases of DNA replication to maintain genome stability.

DNA replication is also regulated in a temporal manner where specific DNA sequences replicate at precise times during S phase, a process known as the DNA replication timing program. While euchromatin replicates in the early part of S phase, heterochromatin and other repressive chromatin types replicate in the later portion of S phase (Gilbert, 2002; Rhind and Gilbert, 2013). Although the process of replication timing has been appreciated for many years, the underlying molecular mechanisms controlling timing have remained elusive. The discovery of factors that regulate the DNA replication timing program, however, demonstrate that replication timing is an actively regulated process.

Once factor that regulates replication timing from yeast to humans is Rif1 (Rap1-interacting factor 1). Rif1 was initially identified as a regulator of telomere length in budding yeast (Hardy et al., 1992), but this function of Rif1 appears to be specific to yeast (Xu, 2004). Subsequently, Rif1 has been shown to regulate multiple aspects of DNA replication and repair. In mammalian cells, Rif1 has been shown to regulate DNA repair pathway choice by preventing resection of double-strand breaks and favoring non-homologous end joining (NHEJ) over homologous recombination (Chapman et al., 2013; Di Virgilio et al., 2013; Zimmermann et al., 2013). Rif1 from multiple organisms contains a Protein Phosphatase 1 (PP1) interaction motif and Rif1 is able to recruit PP1 to DDK-activated helicases to inactive them and prevent initiation of replication (Davé et al., 2014; Hiraga et al., 2014; 2017).

In yeasts, flies and mammalian cells, Rif1 has been shown to regulate the replication timing program (Cornacchia et al., 2012; Hayano et al., 2012; Peace et al., 2014; Sreesankar et al., 2015; Yamazaki et al., 2012). The precise mechanism(s) through which Rif1 functions to control replication timing are not fully understood. For example, Rif1 has been show to interact with Lamin and is thought to tether specific regions of the genome to the nuclear periphery (Foti et al., 2015). How this activity is related to Rif1’s ability to inactivate helicases together with PP1 in controlling the timing program remains obscure.

Studying DNA replication in the context of development provides a powerful method to understand how DNA replication is regulated both spatially and temporally. Although DNA replication is a highly ordered process, it must be flexible enough to accommodate the changes in S phase length and cell cycle parameters that occur as cells differentiate (Matson et al., 2017). For example, during Drosophila development the length of S phase can vary from ∼8 hours in a differentiated mitotic cell to 3-4 minutes during early embryonic cell cycles (Blumenthal et al., 1974; Spradling and Orr-Weaver, 1987). Additionally, many tissues and cell types in Drosophila are polyploid, having multiple copies of the genome in a single cell (Edgar and Orr-Weaver, 2001; Lilly and Duronio, 2005; Zielke et al., 2013).

In polyploid cells, copy number is not always uniform throughout the genome (Rudkin,1969; Hua and Orr-Weaver, 2017; Spradling and Orr-Weaver, 1987). Both heterochromatin and several euchromatic regions of the genome have reduced DNA copy number relative to overall ploidy (Nordman et al., 2011). Underreplicated euchromatic regions of the genome share key features with common fragile sites in that they are devoid of replication origins, late replicating, display DNA damage and are tissue-specific (Andreyeva et al., 2008; Nordman et al., 2014; Sher et al., 2012; Yarosh and Spradling, 2014). The presence of underreplication is conserved in mammalian cells, but the mechanism(s) mammalian cells use to promote underreplication is unknown (Hannibal et al., 2014). In Drosophila, underreplication is an active process that is largely dependent on the Suppressor of Underreplication protein, SUUR (Makunin et al., 2002; Nordman and Orr-Weaver, 2015).

Understanding how the SUUR protein functions will significantly increase our understanding of the developmental control of DNA replication. The SUUR protein has a recognizable SNF2-like chromatin remodeling domain at its N-terminus, but based on sequence analysis, this domain is predicted to be defective for ATP binding and hydrolysis (Makunin et al., 2002; Nordman and Orr-Weaver, 2015). Outside of the SNF2 domain, SUUR has no recognizable motifs or domains, which has hampered a mechanistic understanding of how SUUR promotes underreplication. Recently, however, SUUR was shown to control copy number by directly reducing replication fork progression (Nordman et al., 2014). SUUR associates with active replication replication forks and while loss of SUUR function results in increased replication fork progression, overexpression of SUUR drastically inhibits replication fork progression without affecting origin firing (Nordman et al., 2014; Sher et al., 2012). These findings, together with previous work showing that loss of SUUR function has no influence on ORC binding (Sher et al., 2012) and that SUUR associates with euchromatin in an S phase-dependent manner (Kolesnikova et al., 2013), further supports SUUR as a direct inhibitor of replication fork progression within specific regions of the genome. The mechanism through which SUUR is recruited to replication forks and how it inhibits their progression remains poorly understood.

Here we investigate how SUUR is recruited to replication forks and how it inhibits fork progression. We show that localization of SUUR to replication forks, but not heterochromatin, is dependent on its SNF2 domain. We identify a physical interaction between SUUR and the conserved replication factor Rif1. Importantly, we demonstrate that underreplication is dependent on *Rif1*. Critically, we have shown that Rif1 localizes to replication forks in an SUUR-dependent manner, where it acts downstream of SUUR to control replication fork progression. Our findings provide mechanistic insight into the process of underreplication and define a new function for Rif1 in replication control.

## RESULTS

### The SNF2 domain is essential for SUUR function and replication fork localization

As a first step in understanding the mechanism of SUUR function, we wanted to define how it is localized to replication forks. SUUR has only one conserved domain: a SNF2-like domain in its N-terminal region that is predicted to be defective for ATP binding and hydrolysis (Makunin et al., 2002; Nordman and Orr-Weaver, 2015). To study the function of SUUR’s SNF2 domain, we generated a mutant in which the SNF2 domain was deleted and the resulting mutant protein was expressed under the control of the endogenous *SuUR* promoter. This mutant, *SuUR*^Δ*SNF*^, was then crossed to an *SuUR* null mutant so that it was the only form of the the SUUR protein present. We tested the function of the SuUR^ΔSNF^ mutant protein by assessing its ability to promote underreplication in the larval salivary gland. We purified genomic DNA from larval salivary glands isolated from wandering 3^rd^ instar larvae and generated genome-wide copy number profiles using Illumina-based sequencing. We compared the results we obtained from the *SuUR*^Δ*SNF*^ mutant to copy number profiles from wild-type (WT) and *SuUR* null mutant salivary glands. To identify underreplicated domains, we used CNVnator, which identifies copy number variants (CNVs) based on a statistical analysis of read depth (Abyzov et al., 2011). To be called as underreplicated, regions must not be called as underreplicated in 0-2 hour embryo samples that have uniform copy number and must be larger than 10kb.

The effect of deleting the SNF2 domain was qualitatively and quantitatively similar to the *SuUR* null mutant. Qualitatively, underreplication was suppressed in the *SuUR*^Δ*SNF*^ mutant and the copy number profile was similar to the *SuUR* null mutant (Figure 1B and Supplemental Figure 1). Quantitatively, out of the 90 underreplicated sites identified in WT salivary glands, 59 were not detected in the *SuUR*^Δ*SNF*^ mutant (Supplementary Table 1) and copy number was significantly increased in the euchromatic underreplicated domains similar to the *SuUR* null mutant (Figure 1C). We validated our deep-sequencing findings using quantitative droplet digital PCR (ddPCR) at four underreplicated domains (Figure 1D). Our findings show that the SNF2-like domain of SUUR is necessary to promote underreplication.

**Figure 1.**
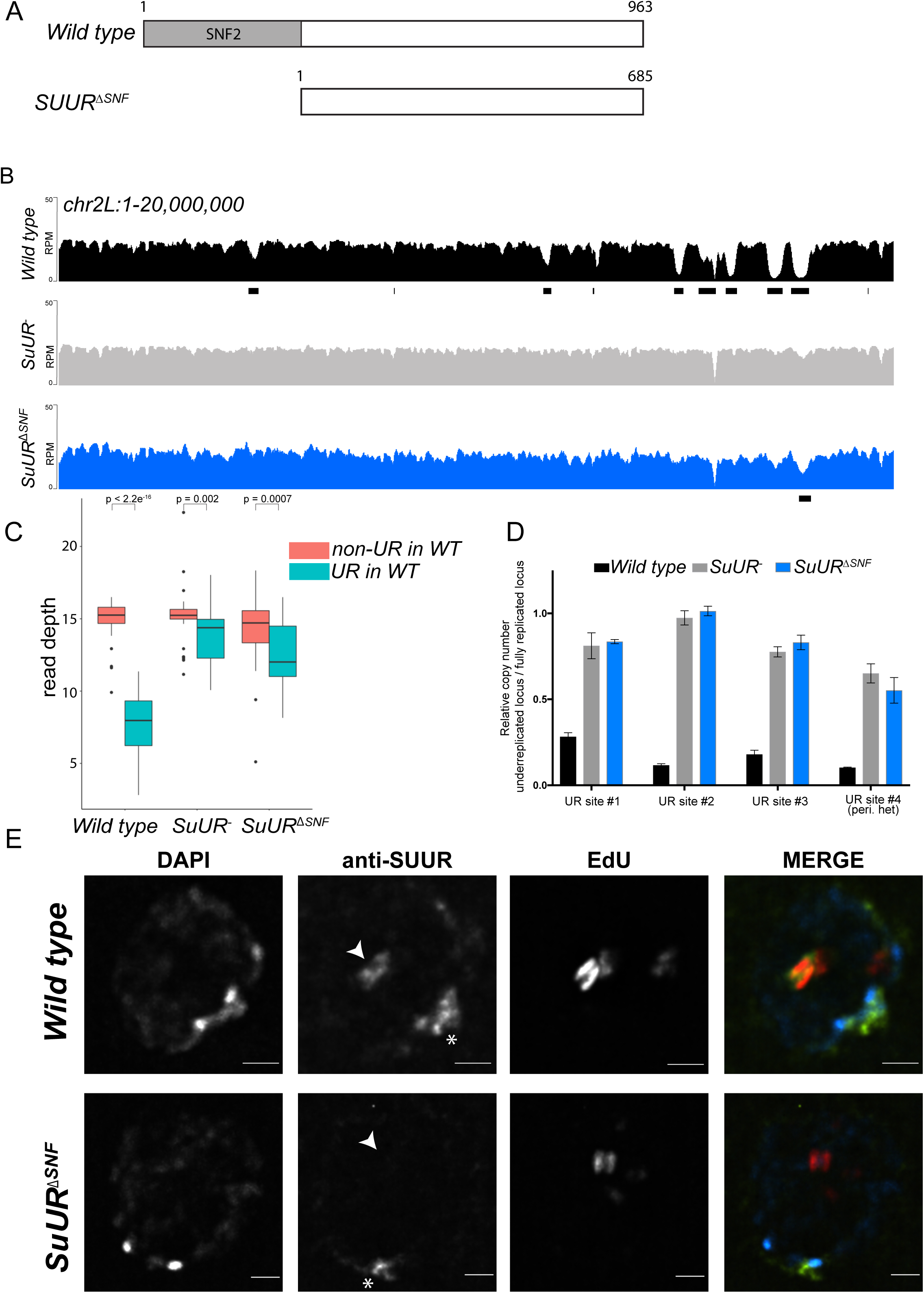
The SNF2 domain is essential for SUUR function and replication fork localization. (A) Schematic representation of the SUUR and SUUR^ΔSNF^ proteins. (B) Illumina-based copy number profiles (Reads Per Million; RPM) of *chr2L* 1-20,000,000 from larval salivary glands. Black bars below each profile represent underreplicated regions identified by CNVnator. (C) Average read depth in regions of euchromatic underreplication domains called in wild-type salivary glands vs. the full replicated regions of the genome. A Welch Two Sample t-test was used to determine *p* values. (D) Quantitative droplet-digital PCR (ddPCR) copy number assay for multiple underreplicated regions. Each bar is the average enrichment relative to fully replicated control region for three biological replicates. Error bars are the SEM. (E) Localization of SUUR in wild-type and *SuUR*^*ΔSNF*^ mutant follicle cells. A single representative stage 13 follicle cell nucleus is shown. Arrowheads indicate sites of amplification. Asterisk marks the chromocenter (heterochromatin). Scale bars are 2μm. DAPI=blue, SUUR=green, EdU=red.

To determine if the SUUR^ΔSNF^ protein was still able to associate with chromatin, we localized SUUR and the SUUR^ΔSNF^ mutant proteins in ovarian follicle cells. During follicle cell development, these cells undergo programmed changes in their cell cycle and DNA replication programs (Claycomb and Orr-Weaver, 2005; Hua and Orr-Weaver, 2017). At a precise time in their differentiation program, follicle cells cease genomic replication and amplify six defined sites of their genome through a re-replication based mechanism. Early in this gene amplification process, both initiation and elongation phases of replication are coupled. Later in the process, however, initiation no longer occurs and active replication forks can be visualized by pulsing amplifying follicle cells with 5-ethynyl-2′deoxyuridine (EdU) (Claycomb et al., 2002). Active replication forks resolve into a double-bar structure, where each bar represents a series of active replication forks travelling away from the origin of replication (Claycomb and Orr-Weaver, 2005). By monitoring SUUR localization in amplifying follicle cells, we can unambiguously determine if SUUR associates with active replication forks.

SUUR has two distinct modes of chromatin association during the endo cycle. It constitutively localizes to heterochromatin and dynamically associates with replication forks (Kolesnikova et al., 2013; Nordman et al., 2014; Swenson et al., 2016). In agreement with previous studies, SUUR localized to both replication forks and heterochromatin in amplifying follicle cells (Figure 1E) (Nordman et al., 2014). In contrast, the SUUR^ΔSNF^ mutant localized to heterochromatin, but its recruitment to active replication forks was severely reduced (Figure 1E). Together, these results demonstrate that the SNF2 domain is important for SUUR recruitment to replication forks and is essential for SUUR-mediated underreplication.

### SUUR associates with Rif1

Interestingly, overexpression of the SNF2 domain and C-terminal portion of SUUR have different underreplication phenotypes. Whereas overexpression of the C-terminal two-thirds of SUUR promotes underreplication (Kolesnikova et al., 2005), overexpression of the SNF2 domain suppresses underreplication in the presence of endogenous SUUR (Kolesnikova et al., 2005). The C-terminal region of SUUR, however, has no detectable homology or conserved domains (Makunin et al., 2002). These observations, together with our own results demonstrating that the SNF2 domain of SUUR is responsible its localization to replication forks, led us to hypothesize that SUUR is recruited to replication forks through its SNF2 domain where it could recruit an additional factor(s) through its C-terminus to inhibit replication fork progression.

To test the hypothesis that a critical factor interacts with the C-terminal region of SUUR to promote underreplication, we used immunoprecipitation mass spectrometry studies to identify SUUR-interacting proteins. We generated flies that expressed FLAG-tagged full length SUUR or the SNF2 domain of SUUR, immunoprecipitated these constructs and identified associated proteins through mass spectrometry. If SUUR recruits a factor to replication forks outside of its SNF2 domain, then we would expect this factor to be present only in full length purifications and not in the SNF2 domain purification. A single protein fulfilled this criteria: Rif1 (Table 1). This result raises the possibility Rif1 works together with SUUR to inhibit replication fork progression.

**Table 1.**

| Table 1 | Full length SUUR |  |  | SNF2 domain |  |  | negative control |  |  |
| --- | --- | --- | --- | --- | --- | --- | --- | --- | --- |
|  | Repl. #1 | Repl. #2 | Repl. #3 | Repl. #1 | Repl. #2 | Repl. #3 | Repl. #1 | Repl. #2 | Repl. #3 |
| SUUR | 36 | 48 | 26 | 21 | 18 | 10 | 1 | 1 | 2 |
| Rif1(CG30085) | 29 | 24 | 14 | 1 | 0 | 0 | 0 | 0 | 0 |

| Comparison of Rif1 abundance* |  |
| --- | --- |
| Full length vs. SNF2 | p < 0.00010 |
| Full length vs. neg. ctrl | p < 0.00010 |

\*Fisher's Exact Test

### Underreplication is dependent on Rif1

If SUUR recruits Rif1 to replication forks to promote underreplication, then underreplication should be dependent on *Rif1*. To test this hypothesis, we used CRISPR-based mutagenesis to generate *Rif1* null mutants in Drosophila (Bassett et al., 2013; Gratz et al., 2013) (Figure 2A). Western blot analysis of ovary extracts from two deletion mutants, *Rif1^1^* and *Rif1^2^,* show no detectable Rif1 protein (Supplemental Figure 2A). Also, no signal was detected in the *Rif1^1^/Rif1^2^* mutant by immunofluorescence (Supplemental Figure 2B). The *Rif1^1^/Rif1^2^* null mutant was viable and fertile showing only a modest defect in embryonic hatch rate relative to wild-type flies with a 92% hatch rate for wild type embryos vs. 88% for the *Rif1^1^/Rif ^2^* mutant embryos (Supplemental Figure 2C). This is in contrast to a previous a study reporting *Rif1* is essential in Drosophila (Sreesankar et al., 2015). Rif1’s essentiality, however, was based on RNAi and not a mutation of the *Rif1* gene (Sreesankar et al., 2015). The most likely explanation for this discrepancy is that the lethality in the RNAi experiments was due to an off-target effect.

**Figure 2.**
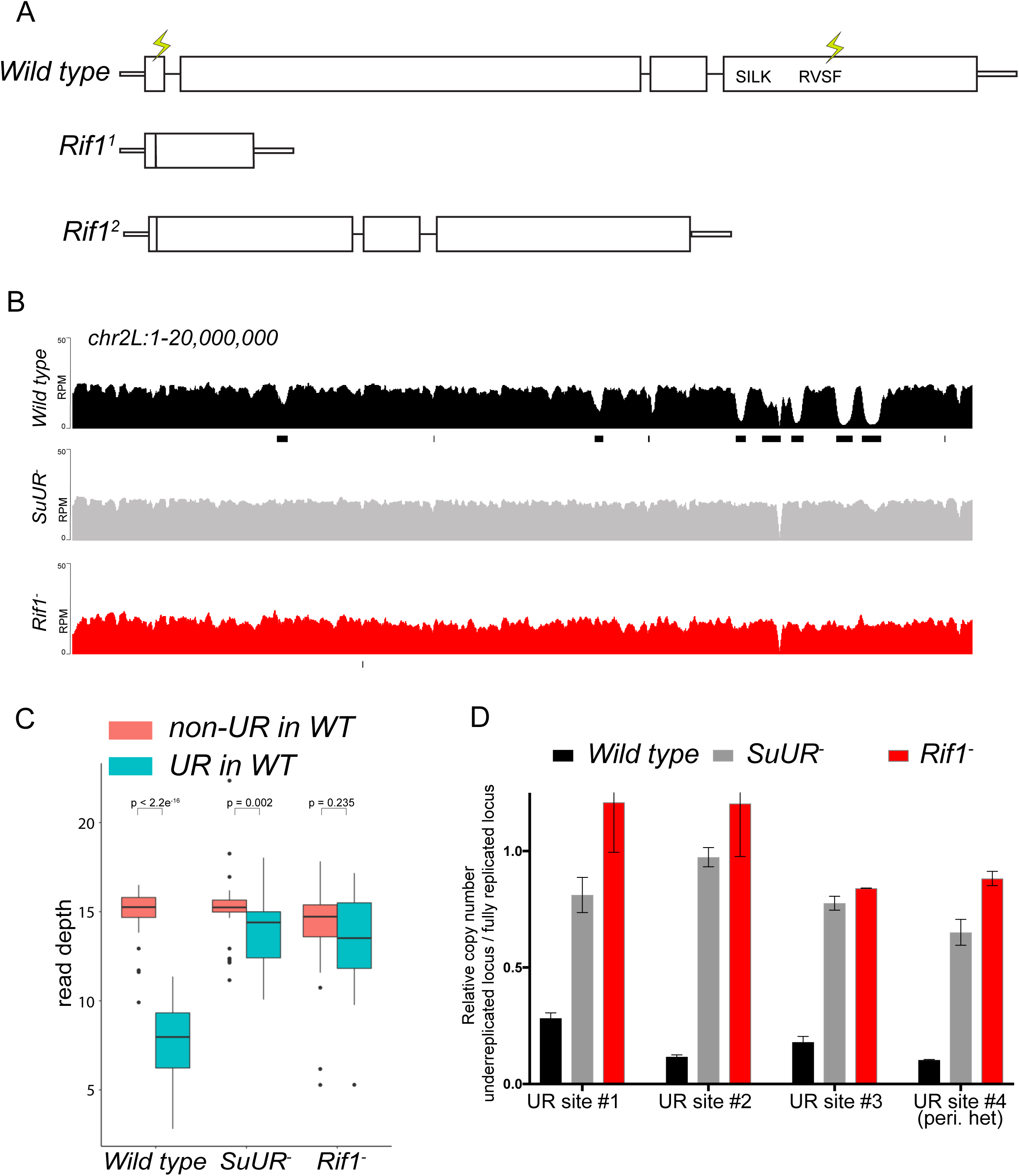
Rif1 is required for underreplication. (A) Schematic representation of the *Rif1* gene and CRISPR-induced *Rif1* mutants. Lightning bolts represent the 5′ and 3′ gRNA positions. (B) Illumina-based copy number profiles of the *chr2L* from larval salivary glands. Black bars below each profile represent underreplicated regions identified by CNVnator. The wild-type and *SuUR* profiles are the same as in Figure 1b. (C) Average read depth in regions of euchromatic underreplication domains called in wild-type salivary glands vs. the fully replicated regions of the genome. A Welch Two Sample t-test was used to determine *p* values. (D) Quantitative droplet-digital PCR (ddPCR) copy number assay for multiple underreplicated regions. Each bar is the average enrichment relative to fully replicated control region for three biological replicates. Error bars are the SEM.

To determine if *Rif1* is necessary for underreplication, we dissected salivary glands from *Rif1^1^*/*Rif1^2^* (herein referred to as *Rif1^−^*) heterozygous larvae and extracted genomic DNA for Illumina-based sequencing to measure changes in DNA copy number. Strikingly, underreplication is abolished upon loss of Rif1 function (Figure 2B and C; Supplemental Figure 3). We validated our sequence-based copy number assays with quantitative PCR at a subset of underreplicated regions using ddPCR (Figure 2D). Furthermore, we determined the read density at all euchromatic sites of underreplication called in our wild-type samples, which quantitatively demonstrates that Rif1 is essential for underreplication (Figure 2C). These results demonstrate that underreplication is dependent on *Rif1*.

It is possible that the *Rif1* mutant indirectly influences underreplication through changes in replication timing. Underreplicated domains, both euchromatic and heterochromatic, tend to be late replicating regions of the genome (Belyaeva et al., 2012; Makunin et al., 2002). Therefore, if these regions replicated earlier in S phase in a *Rif1* mutant, then this change could prevent their underreplication. In fact, SUUR associates with late replicating regions of the genome (Filion et al., 2010; Pindyurin et al., 2007). Due to their large polyploid nature, salivary glands cells cannot be sorted to perform genome-wide replication timing experiments. Because heterochromatin replicates exclusively in late S phase, however, late replication can be visualized when EdU is incorporated exclusively in regions of heterochromatin. To assess if *Rif1* mutants have a clear pattern of late replication in larval salivary glands, we isolated salivary glands from early 3^rd^ instar larvae, which are actively undergoing endo cycles. We pulsed these salivary glands with EdU to visualize sites of replication and co-stained with an anti-HP1 antibody to mark heterochromatin. In wild-type salivary glands, only rarely (1 of 238 EdU^+^ cells; 0.4%) did we detect EdU incorporation in regions of heterochromatin (Supplemental Figure 4). This is consistent with the lack of heterochromatin replication due to underreplication. In contrast, in both *SuUR* and *Rif1* mutants, we could readily detect cells that were solely incorporating EdU within regions of heterochromatin (32 of 327 EdU^+^ cells; 9.8% for *SuUR* and 70 of 385 EdU^+^ cells; 18.2% for *Rif1*) (Supplemental Figure 4). Therefore, we conclude that *Rif1* mutants still have a clear pattern of late replication. Given that heterochromatin underreplication is suppressed in a *Rif1* mutant, although it is still late replicating, indicates that replication timing cannot solely explain the lack of underreplication associated with loss of Rif1 function.

While characterizing Rif1’s role in underreplication and patterns of DNA replication in endo cycling cells, we did observe differences in the heterochromatic regions of *SuUR* and *Rif1* mutants. First, although underreplication is suppressed in both mutants (Figure 2 and Supplemental Figure 3), the chromocenters were abnormally large in *Rif1* mutant relative to an *SuUR* mutant as observed by DAPI staining consistent with the ‘fluffy’ enlarged chromocenters seen in Rif1 mutant mouse cells (Supplemental Figure 4) (Cornacchia et al., 2012). Although, this phenotype was present in all endo cycling cells, it was especially dramatic in the ovarian nurse cells (Supplemental Figure 5). Second, Illumina-based copy number profiles revealed an increase in copy number in some pericentric heterochromatin regions in the *Rif1* mutant relative to the *SuUR* mutant (Supplemental Figure 3). Collectively, these results suggest that heterochromatin is partially, but not fully replicated in *SuUR* mutant endo cycling cells, consistent with previous cytological analysis (Demakova et al., 2007). In contrast, loss of Rif1 function appears to completely restore heterochromatic replication in endo cycling cells.

### Rif1 affects replication fork progression

SUUR-mediated underreplication occurs through inhibition of replication fork progression (Nordman et al., 2014; Sher et al., 2012). If SUUR acts together with Rif1 to promote underreplication, then Rif1 is expected to control replication fork progression. DNA combing assays in human and mouse cells from multiple groups have come to different conclusions as to whether Rif1 affects replication fork progression (Alver et al., 2017; Cornacchia et al., 2012; Hiraga et al., 2017; Yamazaki et al., 2012). Rif1, however, has been shown to be associated with replication forks through nascent chromatin capture, an iPOND-like technique used to isolate proteins associated with active replication forks (Alabert et al., 2014). To determine directly if Rif1 controls replication fork progression, we performed copy number assays on amplifying follicle cells.

Gene amplification in ovarian follicle cells occurs at six discrete sites in the genome through a re-replication based mechanism. Copy number profiling of these amplified domains provides a quantitative assessment of the number of rounds of origin firing and the distance replication forks have travelled during the amplification process, allowing us to disentangle the initiation and elongation phases of DNA replication. To determine if Rif1 affects origin firing and/or replication fork progression, we isolated wild-type and *Rif1* mutant stage 13 egg chambers, which represent the end point of the amplification process, and made quantitative DNA copy number measurements. Loss of Rif1 function resulted in an increase in replication fork progression without significantly affecting copy number at the origin of replication at all sites of amplification (Figure 3A).

**Figure 3.**
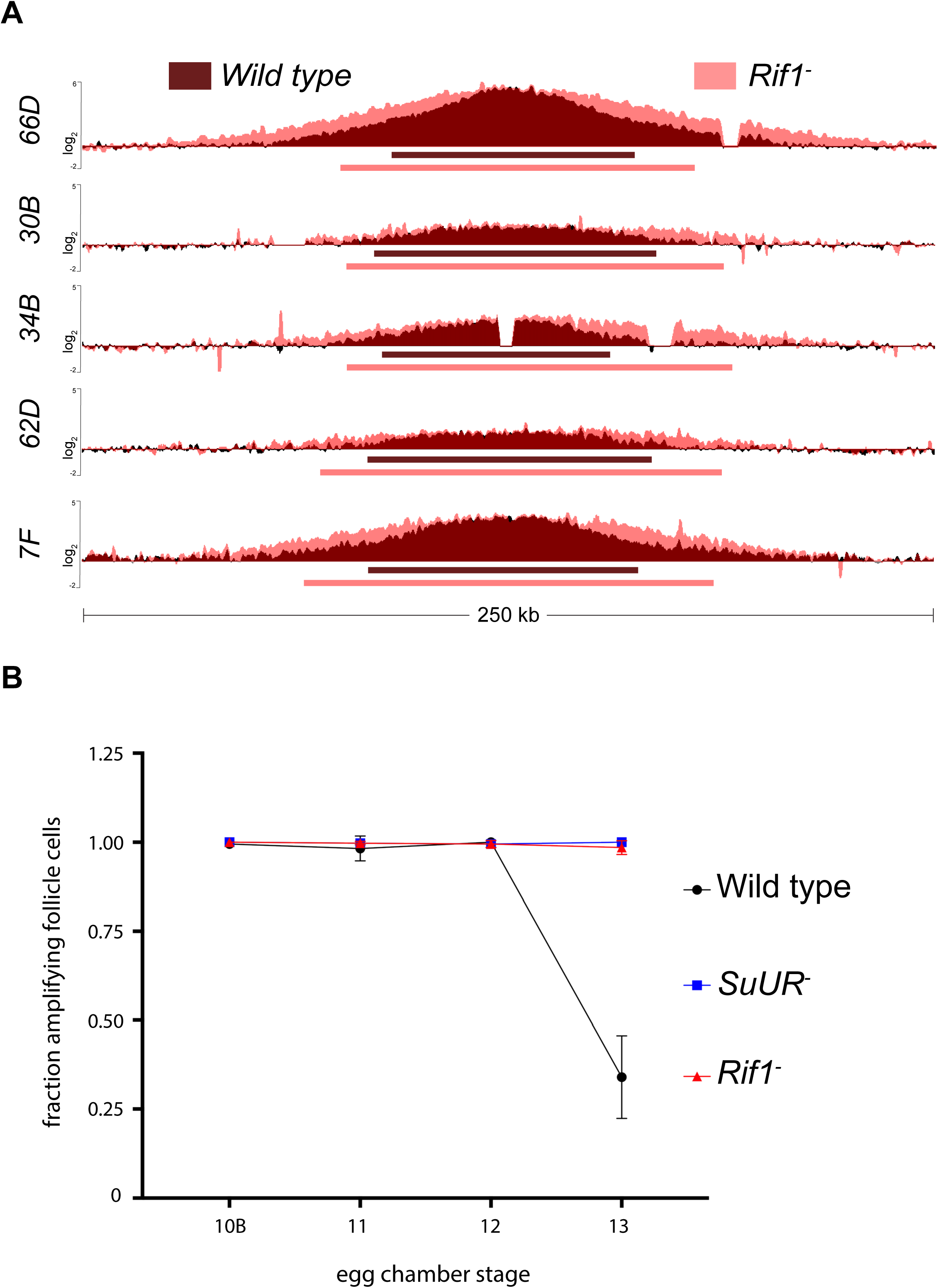
Rif1 regulates replication fork progression. (A) Illumina-based copy number profile of sites of follicle cell gene amplification. DNA was extracted from wild type and *Rif1* mutant stage 13 egg chambers and compared to DNA extracted from 0-2 hr embryos. The resulting graphs are the log2-transformed ratios of egg chamber relative to embryonic DNA. Bars below the graphs represent the distance between the half-maximum copy number on each side of the replication origin. (B) Fraction of cells that display visible amplification foci in each stage of gene amplification. Average of two biological replicates in which two egg chambers from each stage were used per biological replicate. 100- 300 follicle cells were counted per genotype. Error bars are the SEM.

To quantify the changes in fork progression we observed at sites of amplification, we computationally determined the peak of amplification and the region on each arm of the amplified domain that represents one half of the copy number at the highest point of the amplicon (Nordman et al., 2014). This quantitative analysis of origin firing and replication fork progression revealed that origin firing was not affected in the *Rif1* mutant, as no major change in copy number was detected at the origin of replication when comparing wild type and *Rif1* mutant stage 13 follicle cells (Supplemental Table 2). In contrast, the width of each replication gradient, which represents the rate of fork progression, was significantly increased at all sites of amplification (Figure 3A; Supplemental Table 2). Based on the observation that the *Rif1* mutant does not affect origin firing, but specifically affects the distance replication forks travel during the gene amplification process, we conclude that Rif1 regulates replication fork progression.

Given that the *Rif1* mutant phenocopies an *SuUR* mutant with respect to replication fork progression, we next wanted to determine the cause of increased replication fork progression at amplified loci upon loss of Rif1 function. Previously, it was shown that a prolonged period of gene amplification in the *SuUR* mutant gives rise to the extended replication gradient at sites of amplification (Nordman et al., 2014). Gene amplification starts synchronously in all follicle cells at stage 10B of egg chamber development (Calvi et al., 1998). By the end of gene amplification, however, only a subset of follicle cells display visual amplification foci as judged by EdU incorporation (Nordman et al., 2014). To determine if Rif1 controls replication fork progression by increasing the period of gene amplification comparable to an *SuUR* mutant, we quantified the fraction of stage 13 follicle cells that were EdU positive. Similar to an *SuUR* mutant, loss of Rif1 function also resulted in a prolonged period of EdU incorporation with 34% of follicle cells visibly incorporating EdU in wild type follicle cells, 100% in an *SuUR* mutant and 98.5% in the *Rif1* mutant (Figure 3B). This results suggests that Rif1 has a destabilizing effect on replication forks, resulting in a premature cessation of replication fork progression.

### Rif1 acts downstream of SUUR

Rif1 could control SUUR activity and underreplication by at least two different mechanisms. Rif1 could act upstream of SUUR and directly or indirectly regulate SUUR’s ability to associate with chromatin. For example, Histone H1 and HP1 affect underreplication by influencing SUUR’s ability to associate with chromatin (Andreyeva et al., 2017; Pindyurin et al., 2008). Alternatively, Rif1 could act downstream of SUUR to control replication fork progression. We sought to distinguish between these possibilities by determining whether SUUR could still associate with replication forks in the absence of Rif1 function.

To monitor SUUR’s association with heterochromatin and replication forks in the same cell type, we localized SUUR in amplifying follicle cells where replication forks (double bars) and heterochromatin (chromocenter) can be visualized unambiguously, in the presence and absence of Rif1. SUUR localized to both replication forks and heterochromatin in the absence of Rif1 function (Figure 4). Therefore, we conclude that Rif1 acts downstream of SUUR to inhibit fork progression and that SUUR lacks the ability to inhibit replication fork progression in the absence of Rif1.

**Figure 4.**
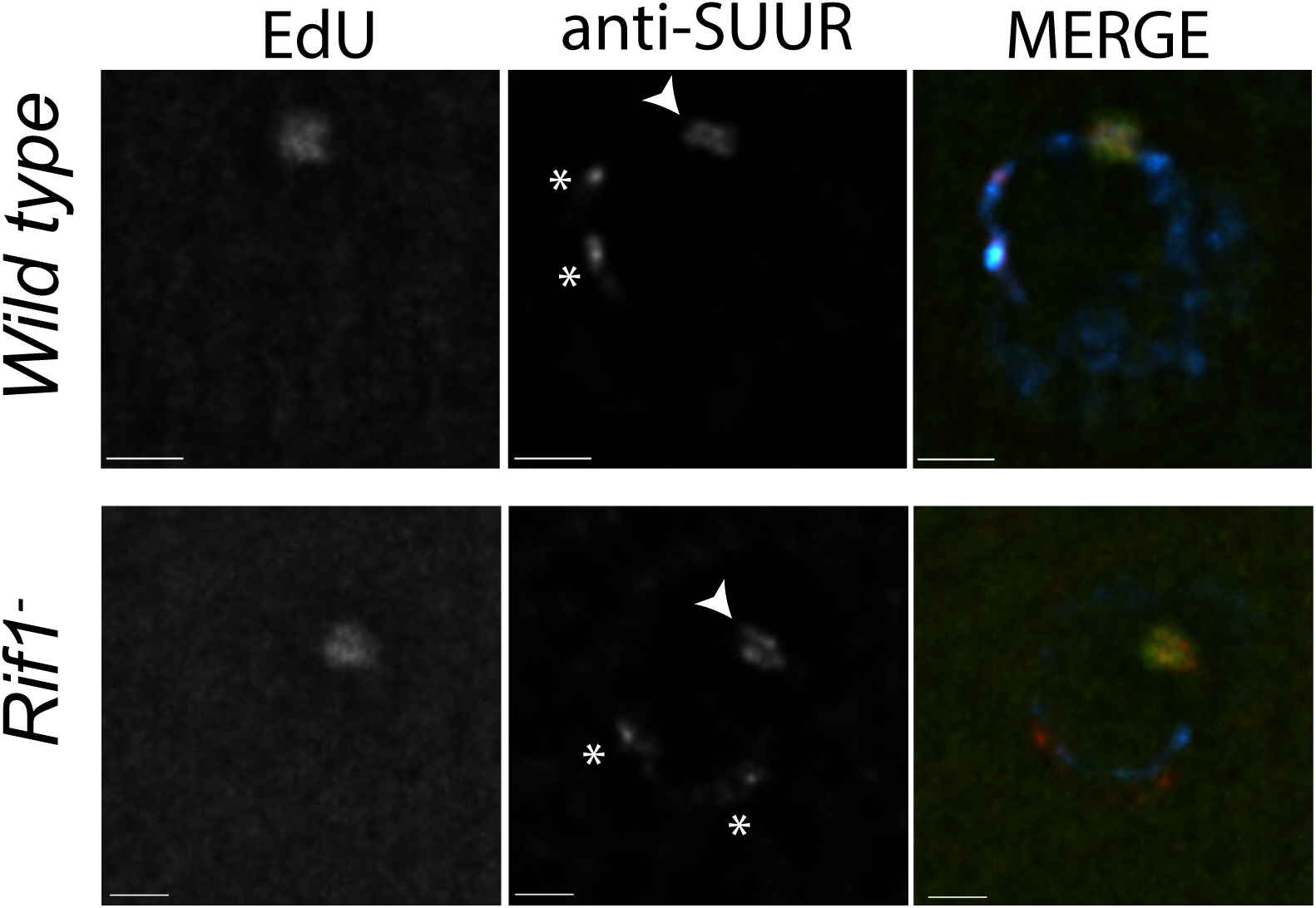
Rif1 acts downstream of SUUR. Localization of replication forks (EdU) and SUUR in a wild-type and *Rif1* mutant follicle cell nuclei. A single representative stage 13 follicle cell nucleus is shown. Scale bars are 2μm. Arrowheads indicate sites of amplification. Asterisks marks the chromocenter (heterochromatin). DAPI=blue, SUUR=green, EdU=red.

### Rif1 localizes to active replication forks

Although our genetic data indicate that Rif1 affects replication fork progression, we wanted to determine if Rif1 controls replication fork progression through a direct or indirect mechanism. If Rif1 directly influences replication fork progression and/or stability, then it should localize to active replication forks. To assess this possibility, we visualized Rif1 localization during gene amplification in follicle cells using a Rif1-specific antibody (Supplemental Figure 2). Rif1 localization pattern was strikingly similar to that of SUUR. First, Rif1 is localized to heterochromatin in all amplification stages amplifying follicle cells (Figure 5). Second, Rif1 localized to sites of amplification even prior to the formation of double bar structures, with weak staining in early stage follicle cells and more intense staining as amplification progressed. Third, in the later stages of gene amplification Rif1 was localized to active replication forks. Taken together, these results demonstrate that Rif1 dynamically associates with the replication forks to regulate their progression.

**Figure 5.**
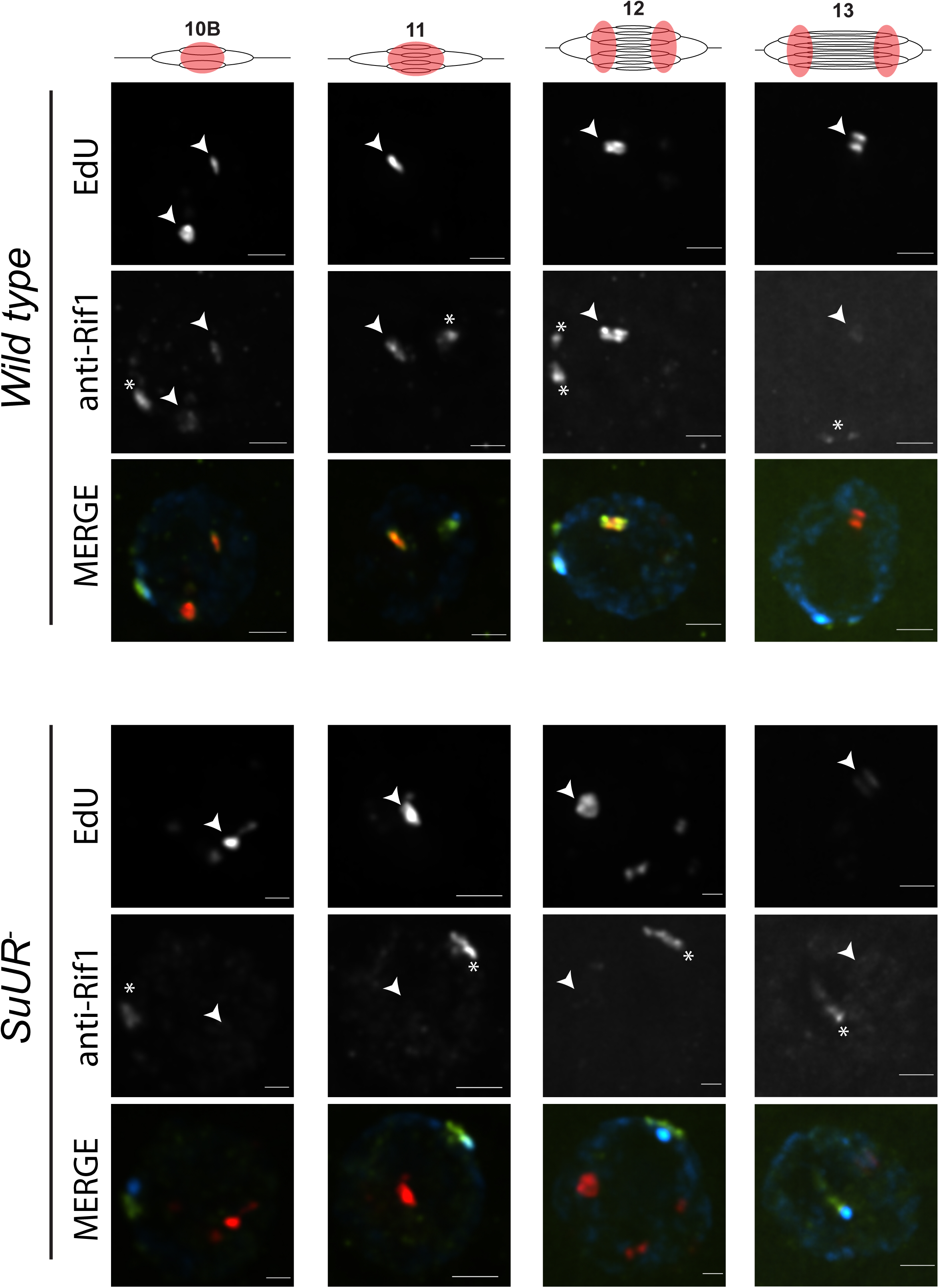
SUUR is necessary to retain Rif1 at replication forks. Localization of active replication forks (EdU) and Rif1 in a wild-type and *SuUR* mutant follicle cell nuclei. Single representative follicle cell nuclei are shown for each stage. Scale bars are 2μm. Arrowheads indicate sites of amplification. Asterisk marks the chromocenter (heterochromatin).

### SUUR is required to retain Rif1 at replication forks

Based on our observations that SUUR physically associates with Rif1 and that a *Rif1* mutant phenocopies an *SuUR* mutant, we hypothesized that SUUR recruits a Rif1/PP1 complex to replication forks. If true, then Rif1 association with replication forks should be at least partially dependent on SUUR. To test this hypothesis, we monitored the localization of Rif1 in *SuUR* mutant amplifying follicle cells. We found that Rif1’s association with replication forks was largely dependent on SUUR, as the Rif1 signal was lost in late stage amplifying follicle cells in an *SuUR* mutant (Figure 5). Rif1’s recruitment to replication foci, however, was not completely dependent on SUUR. In a subset of stage 10B and 11 egg chambers, when both initiation of replication and fork progression are still coupled, we observed Rif1 localization to amplification foci in a subset of follicle cells (data not shown). Rif1 staining was lost, however, in stage 12 and 13 egg chambers. We conclude that while the initial recruitment of Rif1 to sites of amplification is not completely dependent on SUUR, SUUR is necessary to retain Rif1 at replication forks.

### The PP1-interacting motif of Rif1 is necessary for underreplication

Because Rif1 is known to recruit PP1 to replication origins to regulate initiation, this led us to ask if the same interaction between Rif1 and PP1 is important for Rif1’s regulation of replication fork progression. Rif1 associates with Protein Phosphatase 1 (PP1) through a conserved interaction motif, thereby recruiting PP1 to MCM complexes and inactivating them (Davé et al., 2014; Hiraga et al., 2017; 2014). Based on this model of Rif1 function, we wanted to determine if Rif1’s ability to interact with PP1 was necessary for Rif1-mediated underreplication. We used CRISPR-based mutagenesis to mutate the conserved SILK/RSVF PP1 interaction motif to SAAK/RASA. Western blot analysis showed that mutation of the SILK/RSVF motif did not affect protein stability (Supplemental Figure 6). Mutation of this motif has been shown to disrupt the Rif1/PP1 interaction in organisms from yeast to humans (Alver et al., 2017; Davé et al., 2014; Hiraga et al., 2017; 2014; Mattarocci et al., 2014; Sreesankar et al., 2015; Sukackaite et al., 2017). We isolated salivary glands from *Rif1^PP1^* mutant wandering 3^rd^ instar larvae, extracted DNA and measured the copy number of multiple underreplicated domains. Similar to the *Rif1* mutant, underreplication was completely abolished in the *Rif1^PP1^* mutant (Figure 6A). Thus, a Rif1/PP1 complex is necessary to promote underreplication.

**Figure 6.**
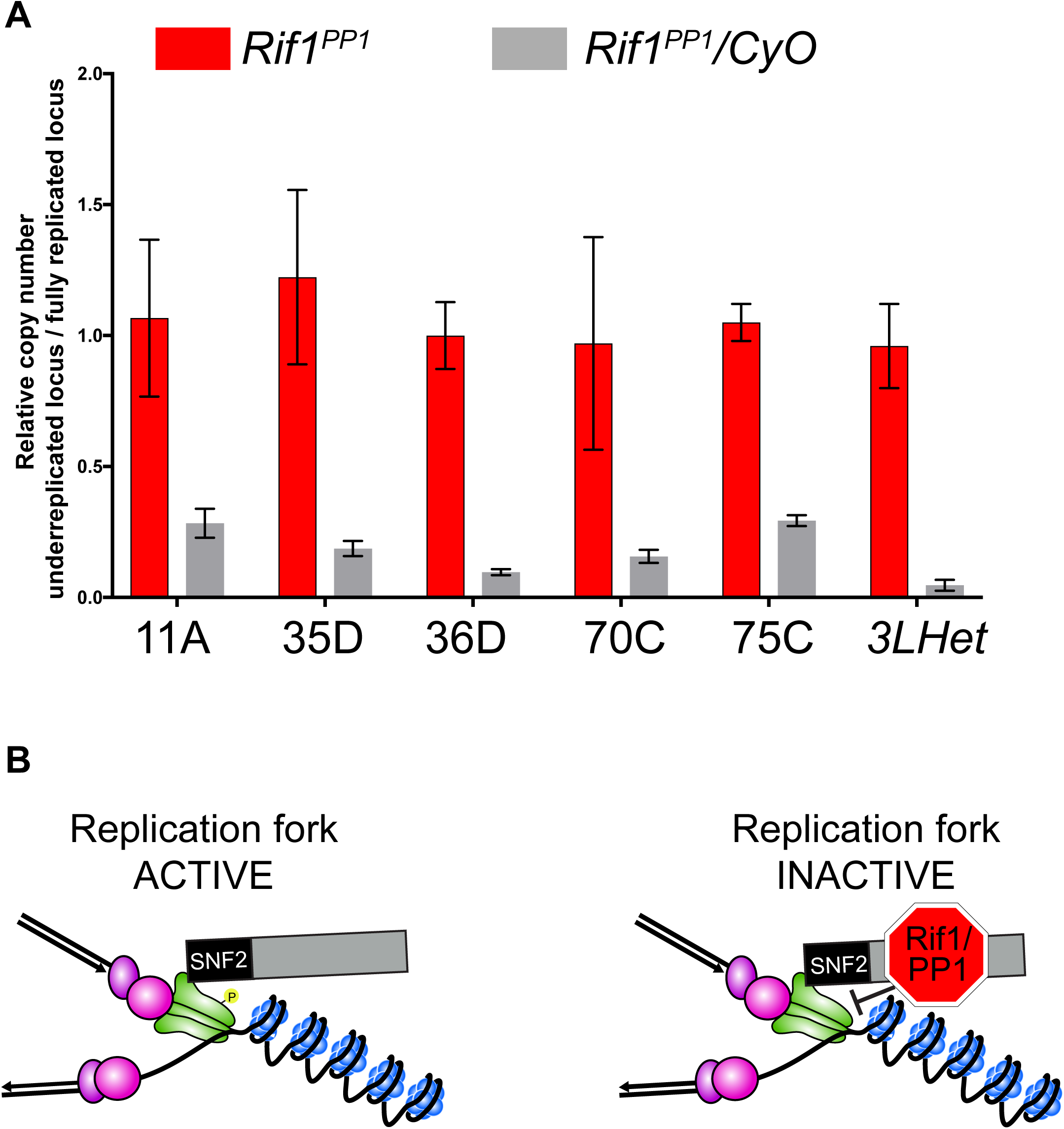
The Rif1/PP1 interaction is necessary to promote underreplication. (A) Quantitative droplet-digital PCR (ddPCR) copy number assay for multiple underreplicated regions. Each bar is the average enrichment relative to fully replicated control region for three biological replicates. Error bars are the SEM. (B) A new model for SUUR-mediated underreplication. In this model SUUR serves as a scaffold to recruit a Rif1/PP1 complex to replication forks where Rif1/PP1 inhibits replication fork progression through dephosphorylation of a component of the replisome. Replication fork image is adapted from (Nordman and Orr-Weaver, 2015)

## DISCUSSION

The SUUR protein is responsible for promoting underreplication of heterochromatin and many euchromatin regions of the genome. Although SUUR was recently shown to promote underreplication through inhibition of replication fork progression, the underlying molecular mechanism has remained unclear. Through biochemical, genetic, genomic and cytological approaches, we have found that SUUR recruits Rif1 to replication forks and that Rif1 is responsible for underreplication. This model is supported by several independent lines of evidence. First, SUUR physically associates with Rif1, and SUUR and Rif1 co-localize at sites of replication. Second, underreplication is dependent on Rif1, although Rif1 mutants have a clear pattern of late replication in endo cycling cells. Third, SUUR localizes to replication forks and heterochromatin in a *Rif1* mutant, however, it is unable to inhibit replication fork progression in the absence of Rif1. Fourth, Rif1 directly controls replication fork progression and phenocopies the effect loss of SUUR function has on replication fork progression. Fifth, SUUR is required for Rif1 localization to replication forks. Critically, using the gene amplification model to separate initiation and and elongation of replication, we have shown that Rif1 can affect fork progression without altering the extent of initiation. Based on these observations, we have defined a new function of Rif1 as a direct regulator of replication fork progression.

### SNF2 domain and fork localization

Our work suggests that the SNF2 domain of SUUR is critical for its ability to localize to replication forks. This is based on the observation that deletion of this domain results in a protein that is unable to localize to replication forks, but still localizes to heterochromatin. SUUR has previously been shown to dynamically localize to replication forks during S phase, but constitutively binds to heterochromatin (Kolesnikova et al., 2013; Nordman et al., 2014). SUUR associates with HP1 and this interaction occurs between the central region of SUUR and HP1. (Pindyurin et al., 2008). Therefore, we speculate that the interaction between SUUR and HP1 is responsible for constitutive SUUR localization to heterochromatin, while a different interaction between the SNF2 domain and a yet to be defined component of the replisome, or replication fork structure itself, recruits SUUR to active replication forks during S phase.

Uncoupling of SUUR’s ability to associate with replication forks and heterochromatin also provides a new level of mechanistic understanding of underreplication. Overexpression of the C-terminal two-thirds of SUUR is capable of inducing ectopic sites of underreplication. In contrast, overexpression of the SUUR’s SNF2 domain, in the presence of endogenous SUUR, suppresses SUUR-mediated underreplication (Kolesnikova et al., 2005). Together with the data presented here, we suggest that overexpression of the SNF2 domain interferes with recruitment of full-length SUUR to replication forks, by saturating potential SUUR binding sites at the replication fork. Although the C-terminal region of SUUR is necessary to induce underreplication (Kolesnikova et al., 2005), the C-terminal portion of SUUR remains associated with heterochromatin in the SuUR^ΔSNF^ construct, but this protein is not sufficient to induce underreplication. We suggest that at physiological levels, the affinity of SUUR with replication forks is substantially diminished in the absence of the SNF2 domain. Our work raises questions about the biological significance of SUUR binding to heterochromatin, since without the SNF2 domain SUUR is still constitutively bound to heterochromatin, yet unable to induce underreplication. Additionally, SUUR dynamically associates with heterochromatin in mitotic cells although heterochromatin is fully replicated (Swenson et al., 2016).

### Rif1 controls underreplication

While trying to uncover the molecular mechanism through which SUUR is able to inhibit replication fork progression, we have uncovered a physical interaction between SUUR and Rif1. Through subsequent analysis, we demonstrated that Rif1 has a direct role in copy number control and that Rif1 acts downstream of SUUR in the underreplication process. Although underreplication is largely dependent on SUUR, there are several sites that display a modest degree of underreplication in the absence of SUUR (Demakova et al., 2007; Sher et al., 2012). In a Rif1 mutant, however, these sites are fully replicated and there is no longer any detectable levels of underreplication within any regions of the genome. It is possible that Rif1 is capable of promoting underreplication through a mechanism independent of SUUR. Therefore, we conclude that Rif1 is a critical factor in driving underreplication.

Further emphasizing the critical role Rif1 plays in copy number control, we have shown that Rif1 acts downstream of SUUR in promoting underreplication. SUUR is still able to associate with chromatin in the absence of Rif1, but is unable to promote underreplication. Underreplicated regions of the genome, including heterochromatin, tend to be late replicating, raising the possibility that changes in replication timing in a *Rif1* mutant suppresses underreplication. *Rif1* mutant endo cycling cells of Drosophila display a cytological pattern of late replication, where heterochromatin is discretely replicated. While Rif1 is likely to control replication timing in Drosophila, we argue that the changes in copy number associated with loss of Rif1 function are not solely due to a loss of late replication. This is supported by the clear pattern of late replication of heterochromatin in *Rif1* mutant endo cycling cells, although heterochromatin appears to be fully replicated in these cells. Previous work in mammalian polyploid cells has shown that underreplication is dependent on Rif1, which was attributed to changes in replication timing (Hannibal and Baker, 2016). It is important to note that Rif1-dependent changes in replication timing were not measured in this system and that many genomic regions transition from early to late replication in a *Rif1* mutant (Foti et al., 2015). Our work raises the possibility that Rif1 has a direct role in mammalian underreplication through a mechanism similar to that of Drosophila and may not simply be due to indirect changes in replication timing. Future work will be necessary to define the role of mammalian Rif1 in underreplication.

### Rif1 regulates replication fork progression

Our analysis of amplification loci demonstrates that Rif1 controls replication fork progression independently of initiation control, thus demonstrating that Rif1 has a specific effect on replication fork progression. Therefore, we have uncovered a new role for Rif1 in DNA metabolism as a regulator of replication fork progression. Rif1 has been identified as part of the replisome in human cells by nascent chromatin capture, a technique that identifies proteins associated with newly synthesized chromatin (Alabert et al., 2014). Multiple studies have assessed whether loss of Rif1 function affects replication fork progression in yeast, mouse and human cells, but have come to different conclusions (Alver et al., 2017; Cornacchia et al., 2012; Hiraga et al., 2017; Yamazaki et al., 2012). DNA fiber assays have been used to measure fork progression in these studies and nearly all have shown that *Rif1* mutants have a slight increase in replication fork progression although not always statistically significant. There could be several reasons for these differing results; Rif1 may control replication fork progression in specific genomic regions that may be underrepresented in some assays, Rif1 function could vary among different cell types, or sample sizes may have been too small to reach significance. Our observations, taken together with these previous studies, leave open the possibility that Rif1-mediated control of replication fork progression could be an evolutionarily conserved function of Rif1. We do not suggest that Rif1 is constitutively associated with replication forks in all cell types. Rather, Rif1 could be recruited to replication forks at a specific time in S phase, or in specific developmental contexts, to modulate the progression of replication forks and provide an additional layer of regulation of the DNA replication program.

How could SUUR and Rif1 function in concert to inhibit replication fork progression? We have shown that Rif1 retention at replication forks is dependent on SUUR. Additionally, underreplication depends on Rif1’s ability to interact with PP1. Rif1/PP1 dephosphorylates DDK-activated helicases to control replication initiation (Davé et al., 2014; Hiraga et al., 2017; 2014). More recently, however, DDK-phosphorylated MCM subunits were shown to be necessary to maintain CMG association and stability of the helicase (Alver et al., 2017). This result suggests that continued phosphorylation of the helicase is necessary for replication fork progression (Alver et al., 2017). We propose that SUUR recruits Rif1/PP1 to replication forks where it is able to dephosphorylate MCM subunits, ultimately inhibiting replication fork progression. Although this mechanism needs to be tested biochemically, it provides a framework to address the underlying molecular mechanism responsible for controlling DNA copy number and could provide new insight into the mechanism(s) Rif1 employs to regulate replication timing.

## MATERIALS AND METHODS

Strain list:

WT – Oregon R
*SuUR^−^* – *w^118^; SuUR^ES^*
*SuUR*^Δ*SNF*^ – *SuUR^ES^*, *PBac*{w^+^ *SuUR^ΔSNF^}*
*Rif1*^−^ – *w^118^*; *Rif1^1^* / *Rif1^2^*
*Rif1^PP1^* – *w^118^*; *Rif1^PP1^*

### BAC-mediated recombineering

BAC-mediated recombineering (Sharan et al., 2009) was used to delete the portion of the *SuUR* gene corresponding to the SNF2 domain. An *attB-P[acman]* clone with a 21-kb genomic region containing the *SuUR* and a *galK* insertion in the *SuUR* coding region (described in (Nordman et al., 2014)) was used as a starting vector. Next, a gene block (IDT) was used to replace the gal*K* cassette and generate a precise deletion within the *SuUR* gene. The resulting vector was verified by fingerprinting, PCR and sequencing. The *SuUR*^Δ*SNF*^ BAC was injected into a strain harboring the *86F8* landing site (Best Gene Inc.).

### Generation of heat shock-inducible, FLAG tagged SuUR transgenic lines

The portion of the *SuUR* gene encoding the SNF2 domain (amino acids 1 to 278) was fused to the SV40 NLS (Barolo et al., 2000) and a 3X-FLAG tag sequence was added to the 5′ end of *SuUR SNF2* sequence. The resulting construct was cloned into the pCaSpeR-hs vector (Thummel and Pirrotta, V.: Drosophila Genomics Resource Center) using the NotI and XbaI restriction sites. A 3X-FLAG tag sequence was added to the 5’ end of of the *SuUR* coding region and cloned into the pCaSpeR-hs vector also using the NotI and XbaI restriction sites. The resulting constructs were verified by sequencing and injected into a *w^1118^* strain (Best Gene Inc.).

### CRISPR mutagenesis

To generate null alleles of *Rif1*, gRNAs targeting the 5′ and 3′ ends of the *Rif1* gene were cloned into the pU6-BbsI plasmid as described (Gratz et al., 2015) using the DRSC Find CRISPRs tool <u>(http://www.flyrnai.org/crispr2/index.html).</u> Both gRNAs were co-injected into a *nos-Cas9* expression stock (Best Gene Inc.). Surviving adults were individually crossed to *CyO*/*Tft* balancer stock and *CyO*-balanced progeny were screened by PCR for a deletion of the *Rif1* locus. Stocks harboring a deletion were further characterized by sequencing. Both *Rif1^1^* and *Rif1^2^* mutants had substantial deletions of the *Rif1* gene and both had frame shift mutations early in the coding region. *Rif1^1^* has a frame shift mutation at amino acid 14, whereas *Rif1^2^* has a frame shift mutation at amino acid 11.

To generate a *Rif1* allele defective for PP1 binding, the pU6-BbsI vector expressing the gRNA targeting the 3′ end of *Rif1* was co-injected with a recovery vector that contained the mutagenized SILK and RVSV (SAAK and RASA) sites with 1kb of homology upstream and downstream of the mutagenized region. Surviving adults were crossed as above and screened by sequencing.

### Cytological analysis and microscopy

Ovaries were dissected from females fattened for two days on wet yeast in Ephrussi Beadle Ringers (EBR) medium (Beadle and Ephrussi, 1935). Ovaries were pulsed with 5-ethynyl-2- deoxyuridine (EdU) for 30 minutes, fixed in 4% formaldehyde and prepared for immunofluorescence (IF) as described (Nordman et al., 2014).

For IF using both anti-Rif1 and anti-SUUR antibodies, ovaries were dissected, pulsed with 50μM EdU and fixed. Ovaries were then incubated in primary antibody (1:200) overnight at 4°C. Alexa Fluor secondary antibodies (ThermoFisher) were used at a dilution of 1:500 for 2 hours at room <u>temperature. EdU</u> detection was performed after incubation of the secondary antibody using Click-iT Alexa Fluor-555 or −488 (Invitrogen). All images were obtained using a Nikon Ti-E inverted microscope with a Zyla sCMOS digital camera. Images were deconvolved and processed using NIS-Elements software (Nikon).

For salivary gland IF,^3rd^ instar larvae were collected prior to the wandering stage. Salivary glands were dissected in EBR, pulsed with 50μM EdU for 30 minutes and fixed with 4% formaldehyde. Salivary glands were incubated in anti-HP1 antibody (Developmental Studies Hybridoma Bank; C1A9) overnight at 4°C. Alexa Fluor secondary antibodies staining and Click-iT EdU labeling were performed as described above.

### Rif1 antibody production

Rif1 antiserum was produced in guinea pigs and rabbits (Cocalico Biologicals Inc.). Briefly, a Rif1 protein fragment from residues 694-1094 (Sreesankar et al., 2012) was C-terminally six-histidine tagged and and expressed in *E. coli* Rossetta DE3 cells and purified using Ni-NTA Agarose beads (Qiagen). The purified protein was used for injection (Cocalico Biologicals Inc.) and serum was affinity purified as described (Moore and Orr-Weaver, 1998). Affinity purified guinea pig anti-Rif1 antibody was used for immunofluorescence.

### IP-mass spec

Flies containing heat shock-inducible *SuUR* transgenes were expanded into population cages. 0- 24 hour embryos were collected, incubated at 37°C for one hour, and allowed to recover for one hour following heat shock treatment. Wild-type embryos were used as a negative control. Embryos were dechorionated in bleach and fixed for 20 minutes in 2% formaldehyde. Approximately 0.5g of fixed and dechorionated embryos were used for each replicate. Embryos were disrupted by douncing in Buffer 1 (Shao et al., 1999), followed by centrifugation at 3,000 × g for 2 minutes at 4°C and resuspended in lysis buffer 3 (MacAlpine et al., 2010). Chromatin was prepared by sonicating nuclei for a total of 40 cycles of 30″ ON and 30″ OFF at max power using a Bioruptor 300 (Diagnenode) with vortexing and pausing after every 10 cycles. Cleared lysates were incubated with anti-FLAG M2 affinity gel (Sigma) for 2 hours at 4°C. After extensive washing in LB3 and LB3 with 1M NaCl, proteins were eluted using 3X FLAG peptide (Sigma). Crosslinks were reversed by boiling purified material in Laemmli buffer with β-mercaptoethanol for 20 minutes.

Immunoprecipitated samples were separated on a 4-12% NuPAGE Bis-Tris gel (Invitrogen), proteins were stained with Novex colloidal Coomassie stain (Invitrogen), and destained in water. Coomassie stained gel regions were cut from the gel and diced into 1mm^3^ cubes. Proteins were reduced and alkylated, destained with 50% MeCN in 25mM ammonium bicarbonate, and in-gel digested with trypsin (10ng/uL) in 25mM ammonium bicarbonate overnight at 37°C. Peptides were extracted by gel dehydration with 60% MeCN, 0.1% TFA, the extracts were dried by speed vac centrifugation, and reconstituted in 0.1% formic acid. Peptides were analyzed by LC-coupled tandem mass spectrometry (LC-MS/MS). An analytical column was packed with 20cm of C18 reverse phase material (Jupiter, 3 μm beads, 300Å, Phenomenox) directly into a laser-pulled emitter tip. Peptides were loaded on the capillary reverse phase analytical column (360 μm O.D. × 100 μm I.D.) using a Dionex Ultimate 3000 nanoLC and autosampler. The mobile phase solvents consisted of 0.1% formic acid, 99.9% water (solvent A) and 0.1% formic acid, 99.9% acetonitrile (solvent B). Peptides were gradient-eluted at a flow rate of 350 nL/min, using a 120-minute gradient. The gradient consisted of the following: 1-3min, 2% B (sample loading from autosampler); 3-98 min, 2-45% B; 98-105 min, 45- 90% B; 105-107 min, 90% B; 107-110 min, 90-2% B; 110-120 min (column re-equilibration), 2% B. A Q Exactive HF mass spectrometer (Thermo Scientific), equipped with a nanoelectrospray ionization source, was used to mass analyze the eluting peptides using a data-dependent method. The instrument method consisted of MS1 using an MS AGC target value of 3e6, followed by up to 15 MS/MS scans of the most abundant ions detected in the preceding MS scan. A maximum MS/MS ion time of 40 ms was used with a MS2 AGC target of 1e5. Dynamic exclusion was set to 20s, HCD collision energy was set to 27 nce, and peptide match and isotope exclusion were enabled. For identification of peptides, tandem mass spectra were searched with Sequest (Thermo Fisher Scientific) against a *Drosophila melanogaster* database created from the UniprotKB protein database (www.uniprot.org). Search results were assembled using Scaffold 4.3.4 (Proteome Software).

### Genome-wide copy number profiling

Embryos were collected immediately after 2 hours of egg laying. Salivary glands were dissected in EBR from 50 wandering 3^rd^ instar larvae per genotype and flash frozen. Ovaries were dissected from females fattened for two days on wet yeast in EBR and 50 stage 13 egg chambers were isolated for each genotype and flash frozen. Tissues were thawed on ice, resuspended in LB3 and dounced using a Kontes B-type pestle. Dounced homogenates were sonicated using a Bioruptor 300 (Diagenode) for 10 cycles of 30″ on and 30″ off at maximal power. Lysates were treated with RNase and Proteinase K and genomic DNA was isolated by phenol-chloroform extraction. Illumina libraries were prepared using NEB DNA Ultra II (New England Biolabs) following the manufacturers protocol. Barcoded libraries were sequenced using Illumina NextSeq500 platform.

### Bioinformatics

Reads were mapped to the Drosophila genome (BDGP Release 6) using BWA using default parameters (Li and Durbin, 2009). CNVnator 0.3.3 was used for the detection of underreplicated regions using a bin size of 1000 (Abyzov et al., 2011). Regions were identified as underreplicated if they were identified as underreplicated in 0-2h embryonic DNA and were greater than 10kb in length. The number of reads for underreplicated regions was called by using bedtools multicov tool for the underreplicated and uncalled regions. Average read depth per region was determined by multiplying the number of reads in a region by the read length and dividing by the total region length. Read depth was normalized between samples by scaling the total reads obtained per sample. Statistical comparison between the regions was with a t-test. For read depth in pericentric heterochromatin regions, the chromatin arm was binned into 10kb windows and the number of reads for each window was called using bedtools multicov using only uniquely mapped reads.

Half maximum analysis of amplicon copy number profiles was performed as described previously (Alexander et al., 2015; Nordman et al., 2014). Briefly, log_2_ ratios were generated using bamCompare from deepTool 2.5.0 by comparing stage 13 follicle cell profiles to a 0-2h embryo sample. Smoothed log_2_-transformed data was used to determine the point of maximum copy number associated with each amplicon. The chromosome coordinate corresponding to half the maximum value for each arm of the amplicon was then determined.

### Copy number analysis by droplet-digital PCR (ddPCR)

Genomic DNA was extracted from salivary glands isolated from wandering 3^rd^ instar larvae as described above. Primer sets annealing to the mid-point of the indicated UR regions were used (previously described in (Nordman et al., 2014; Sher et al., 2012)). ddPCR was performed according to manufacture’s recommendations (BioRad). All ddPCR reactions were performed in triplicate from three independent biological replicates. The concentration value for each set of primers in an underreplicated domain was divided by the concentration value of a fully replicated control to generate the bar graph. Error bars represent the SEM.

### Western blotting

Ovaries were dissected from females fattened for two days on wet yeast and suspended in Laemmli buffer supplemented with DTT. Ovaries were homogenized and boiled and extracts were loaded on a 4-20% Mini-PROTEAN TGX Stain-Free gel (BioRad). After electrophoresis the gel was activated and imaged according to the manufacturers recommendations. Protein was transferred to a PDVF membrane using a Trans-Blot Turbo Transfer System (BioRad). After blocking and incubation with antibodies, blots were imaged using an Amersham 600 CCD imager.

## DATA ACCESS

Data sets described in this manuscript can be found under the GEO accession number: GSE114370.

## ACKNOWLEDGMENTS

We thank Kristie Rose at the Vanderbilt Proteomics core for mass spectrometry and Olivia Koues from the VANTAGE core at Vanderbilt for Illumina sequencing. Terry Orr-Weaver, Stephen Bell, Katherine Friedman, James Dewar, Dave Cortez and members of the Nordman lab for providing critical comments on the manuscript. We thank Brooke Hamilton for assistance in generating the *Rif1* mutants. This work was supported by an NIH R00 award 5R00GM104151 to J.T.N.

## DISCLOSURE

The authors have no conflicts of interest

---

**Supplemental Figure S1 – related to Figure 1.**
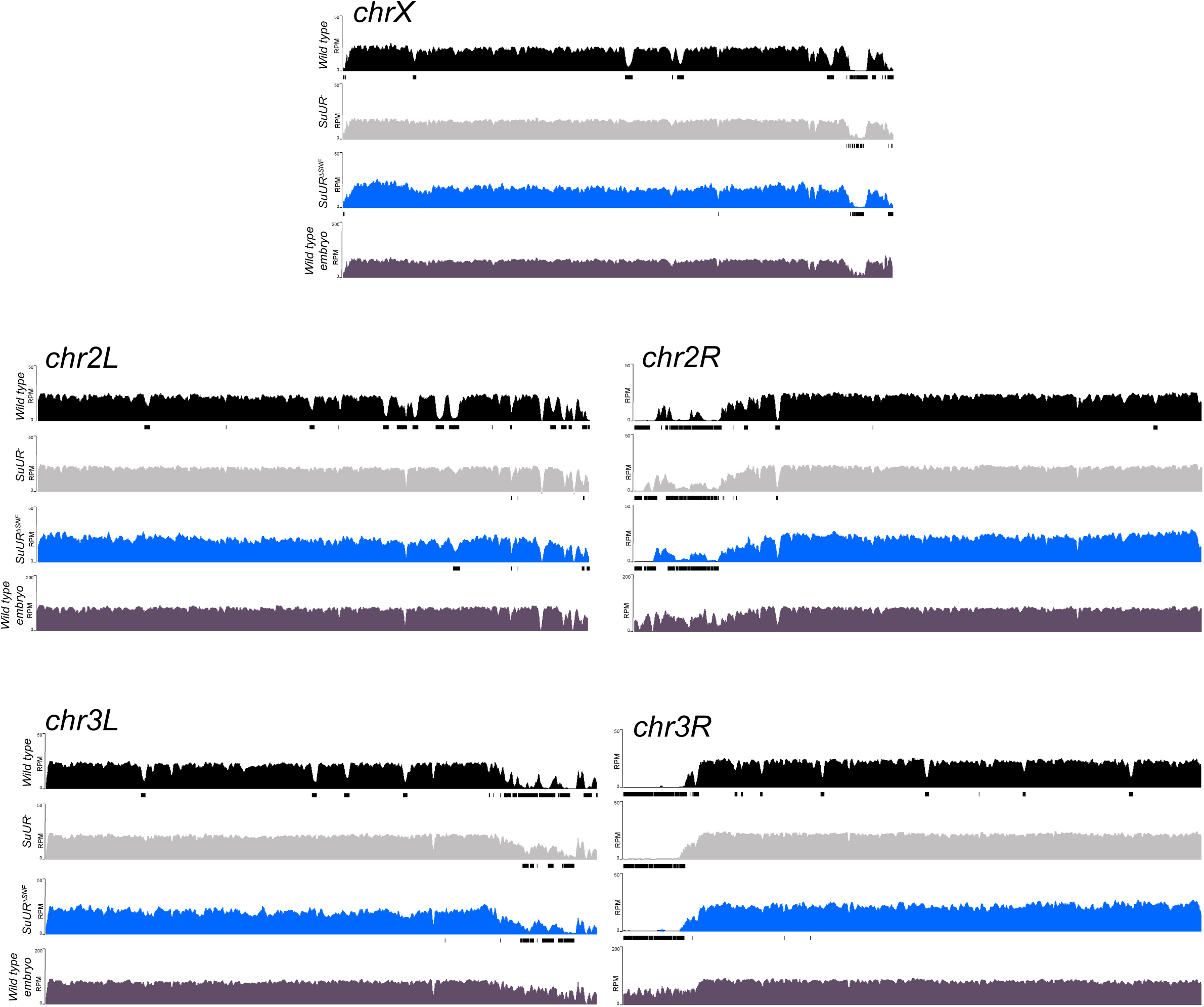
Genome-wide copy number profile of the *SuUR*^*ΔSNF*^ mutant. Illumina-based copy number profiles of all chromosome arms except the fourth for larval salivary glands of the indicated genotypes and wild type 0-2h embryos in which DNA is fully replicated. Black bars below each profile represent called underreplicated regions.

**Supplemental Figure S2 – related to Figure 2 and Figure 5.**
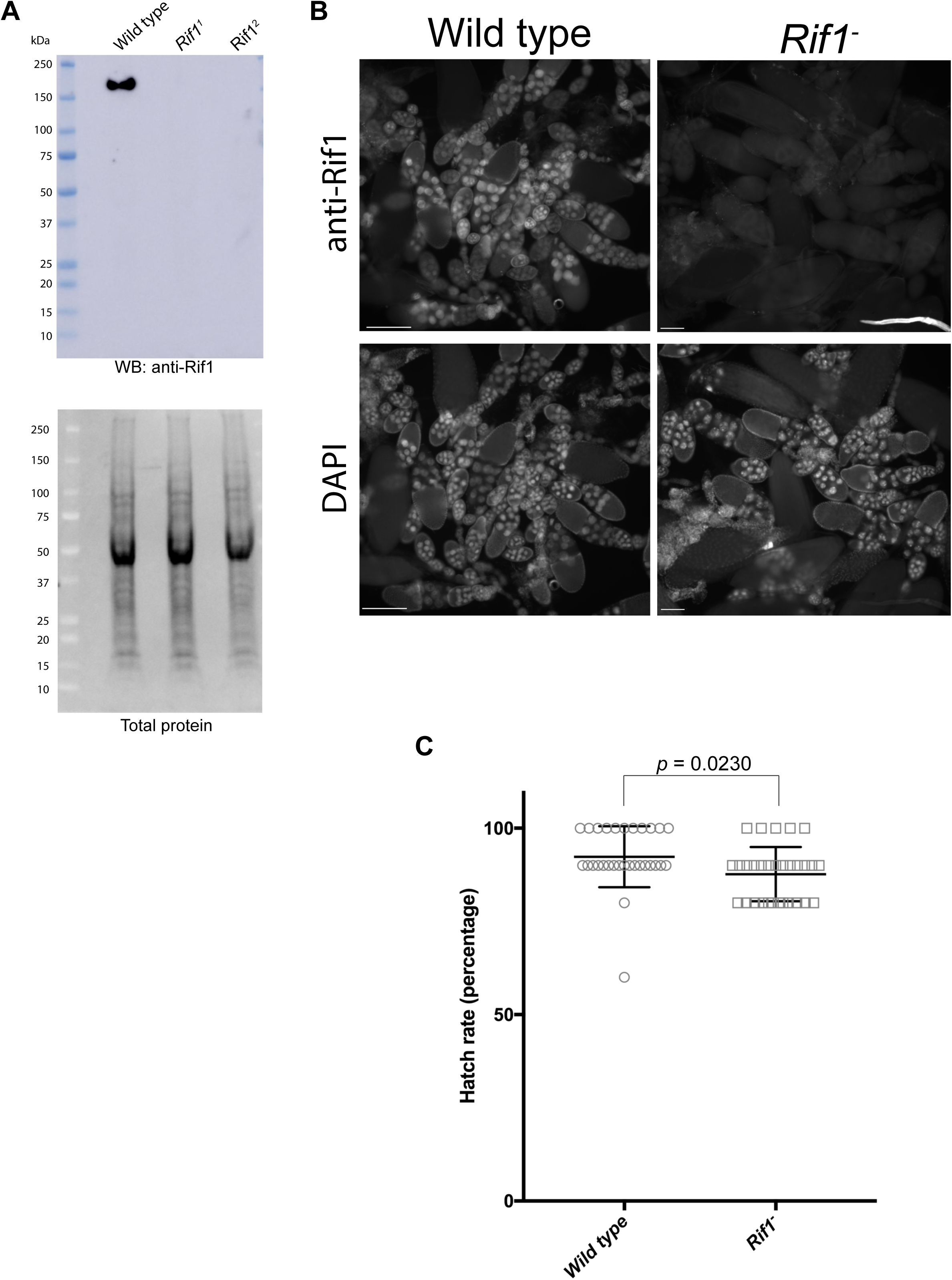
Verification of *Rif1* mutants and validation of anti-Rif1 antibody. (A) Western blot analysis of ovary extracts prepared from the indicated genotypes. Serum produced in guinea pigs was used at 1:1000 dilution. (B) Immunofluorescence of ovaries using affinity purified anti-Rif1 antibody produced in guinea pigs. Exposure times were equal between the two genotypes. (C) Embryo hatch rate assay comparing embryos laid by wild-type or *Rif1^1^/Rif1^2^* mutant mothers. n=300 embryos per genotype. Each data point represents the hatch rate of a group of 10 embryos. An unpaired student t-test was used to generate the *p* value.

**Supplemental Figure S3 – related to Figure 2.**
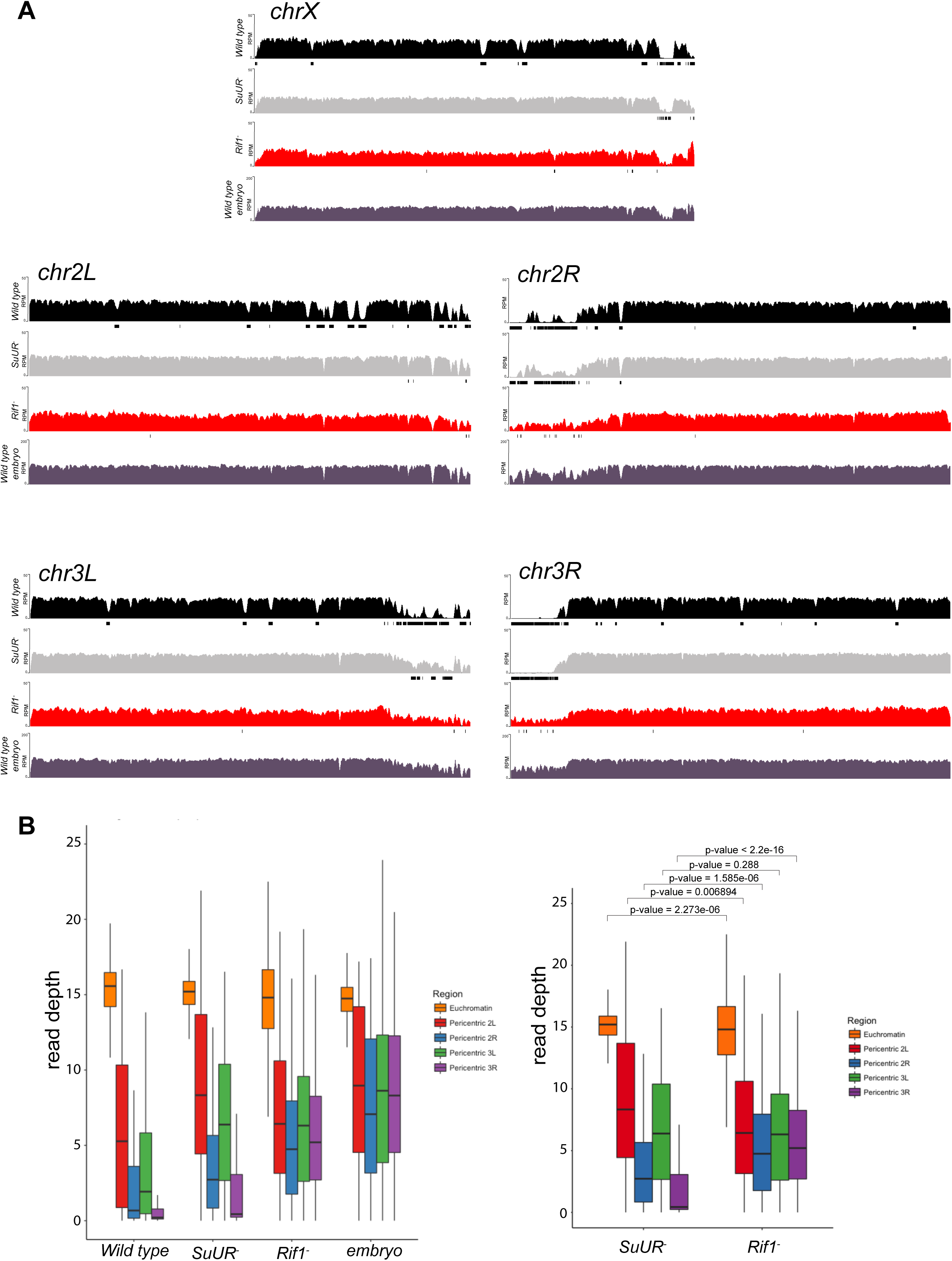
Genome-wide copy number profile of the *Rif1* mutant. (A) Illumina-based copy number profiles of all chromosome arms except the fourth for larval salivary glands of the indicated genotypes. Black bars below each profile represent called underreplicated regions. (B) Box plot represents read depth in 10 kb bins in the pericentric chromatin regions for *chr 2L, 2R, 3L* and *3R*. A Welch Two Sample t-test was used to compare the same regions between *SuUR* and *Rif1* mutants. The same wild-type, *SuUR* and 0-2h embryo plots as in Supplemental Figure S1.

**Supplemental Figure S4.**
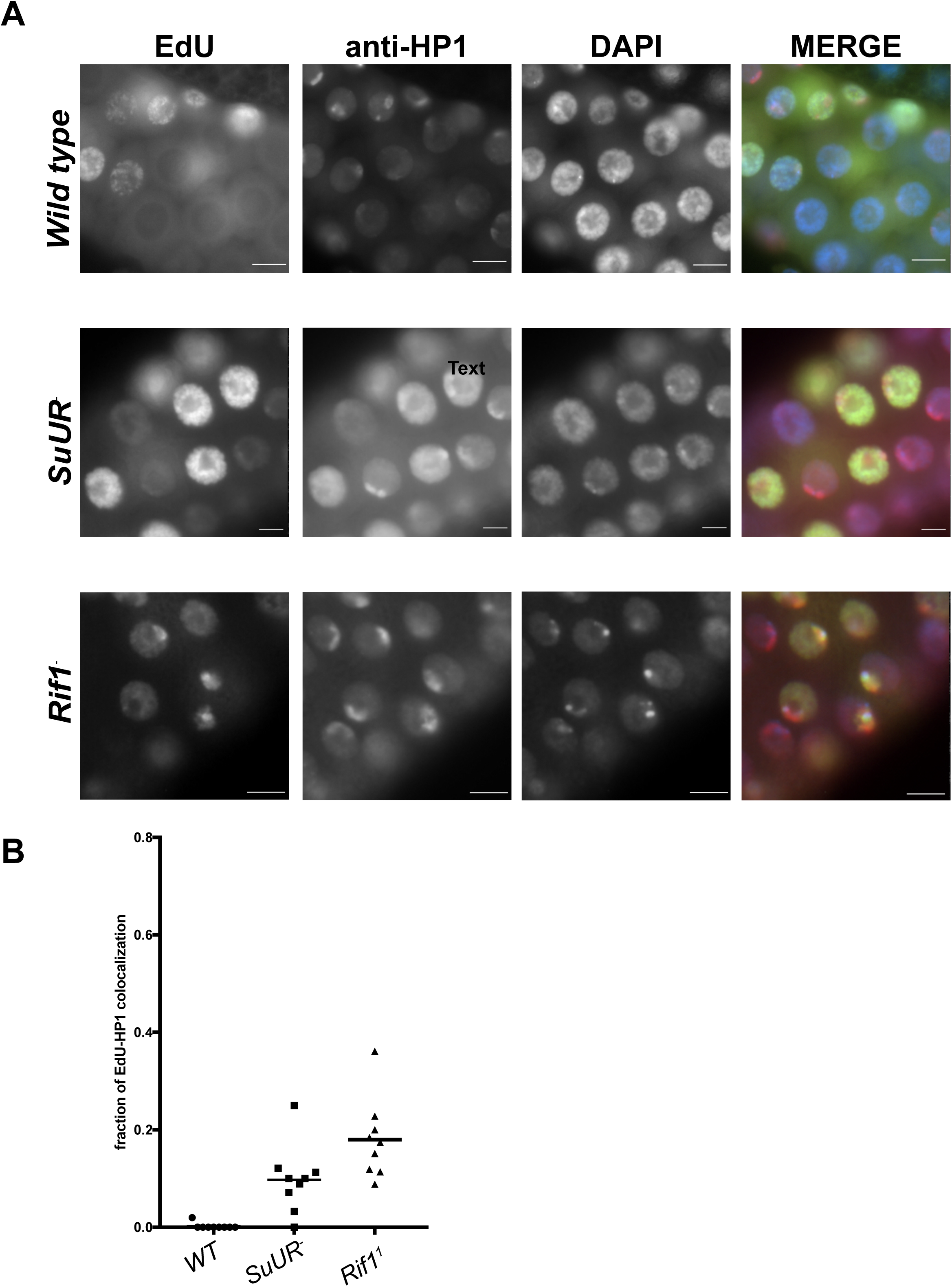
Rif1 mutant salivary gland cells display a pattern of late replication. (A) Representative immunofluorescent images of 3^rd^ instar salivary glands pulse labelled with EdU and stained with anti-HP1 to mark heterochromatin. Wild-type cells fail to incorporate EdU into regions of heterochromatin due to underreplication, whereas EdU can be detected in the heterochromatic regions of *SuUR* and *Rif1* mutants. DAPI=blue, EdU=green, HP1=red (B) Quantitation of three biological replicates. Out of the total number of EdU positive cells, the fraction incorporating EdU predominantly in the heterochromatic (HP1) regions were measured. More than 200 EdU positive cells were scored for each genotype.

**Supplemental Figure S5.**
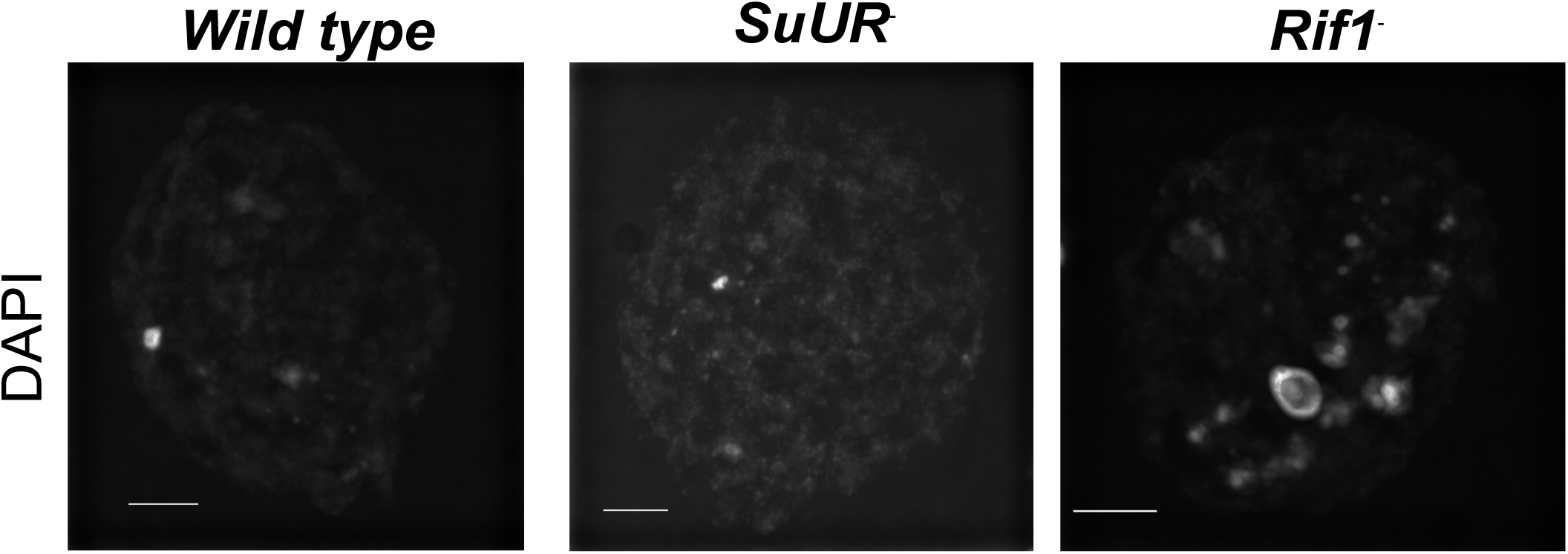
*Rif1* mutant endo cycling cells have enlarged chromocenters. Representative image of of nurse cell nuclei from stage 10 egg chambers. Egg chambers were stained with DAPI. Scale bar is 10 μm. Exposure times and scaling are equal in all images.

**Supplemental Figure S6 – related to Figure 6.**
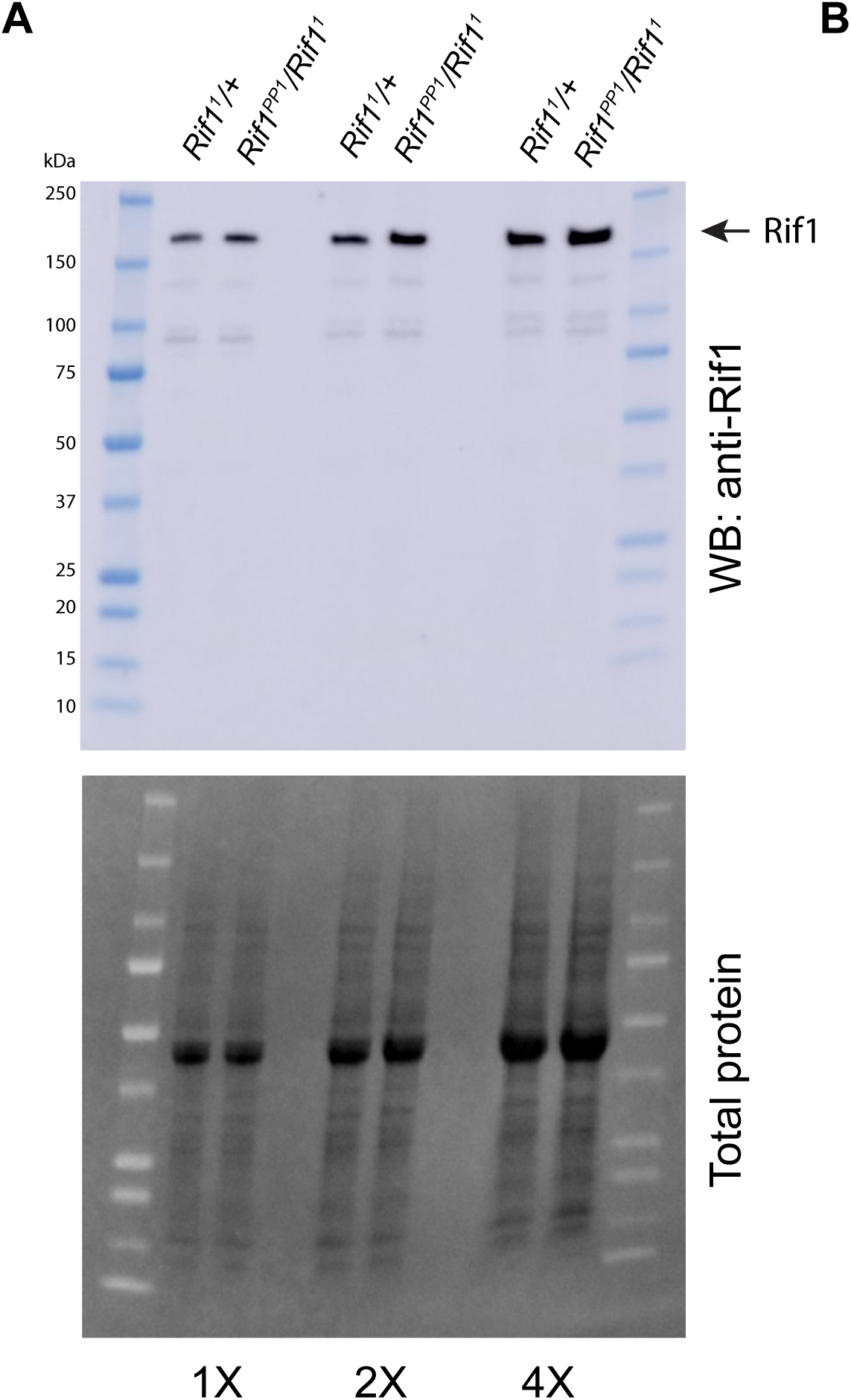
The Rif1^PP1^ protein expression is similar to wild type Rif1. (A) Western blot analysis of ovary extracts from *Rif1^PP1^*/*Rif1^1^* and *Rif1^1^*/*+* adults. Serum was produced in guinea pigs and used at 1:1000 dilution.

**Supplemental Table 1:** Underreplicated regions called by CNVnator

| chr | start | end | OregonR | SuUR- | SuURΔSNF | Rif1- |
| --- | --- | --- | --- | --- | --- | --- |
| chrX | 9001 | 52000 | + |  | + |  |
| chrX | 80001 | 108000 | + |  |  |  |
| chrX | 2976001 | 3125000 | + |  |  |  |
| chrX | 12052001 | 12382000 | + |  |  |  |
| chrX | 14062001 | 14101000 | + |  |  |  |
| chrX | 14292001 | 14563000 | + |  |  |  |
| chrX | 20686001 | 20957000 | + |  |  |  |
| chrX | 20962001 | 20998000 | + |  |  |  |
| chrX | 21525001 | 21544000 | + | + |  | + |
| chrX | 21667001 | 21807000 | + | + | + |  |
| chrX | 21814001 | 21829000 | + |  |  |  |
| chrX | 21843001 | 21895000 | + | + | + |  |
| chrX | 21904001 | 21965000 | + | + | + |  |
| chrX | 21972001 | 22433000 | + | + | + |  |
| chrX | 22611001 | 22787000 | + | + |  |  |
| chrX | 23053001 | 23064000 | + | + |  |  |
| chrX | 23175001 | 23205000 | + |  |  |  |
| chrX | 23285001 | 23535000 | + | + | + |  |
| chr2L | 4538001 | 4777000 | + |  |  |  |
| chr2L | 8013001 | 8024000 | + |  |  |  |
| chr2L | 11587001 | 11777000 | + |  |  |  |
| chr2L | 12781001 | 12799000 | + |  |  |  |
| chr2L | 14722001 | 14946000 | + |  |  |  |
| chr2L | 15306001 | 15721000 | + |  |  |  |
| chr2L | 15960001 | 16154000 | + |  |  |  |
| chr2L | 16161001 | 16217000 | + |  |  |  |
| chr2L | 16948001 | 17199000 | + |  |  |  |
| chr2L | 17201001 | 17312000 | + |  |  |  |
| chr2L | 17520001 | 17951000 | + |  | + |  |
| chr2L | 19345001 | 19358000 | + |  |  |  |
| chr2L | 20128001 | 20199000 | + | + | + |  |
| chr2L | 21833001 | 22085000 | + |  |  |  |
| chr2L | 22284001 | 22522000 | + |  |  |  |
| chr2L | 22614001 | 22752000 | + |  |  |  |
| chr2L | 23198001 | 23406000 | + | + | + | + |
| chr2L | 23415001 | 23513712 | + |  | + | + |
| chr2R | 1 | 699000 | + | + | + | + |
| chr2R | 1194001 | 1206000 | + |  |  |  |
| chr2R | 1379001 | 1501000 | + | + | + |  |
| chr2R | 1557001 | 1589000 | + |  |  |  |

Supplementary Table 1 cont.
|  |  |  |  |  |  |  |
| --- | --- | --- | --- | --- | --- | --- |
| chr2R | 1609001 | 1700000 | + |  |  |  |
| chr2R | 1707001 | 1968000 | + | + | + |  |
| chr2R | 1975001 | 2163000 | + | + | + | + |
| chr2R | 2170001 | 2409000 | + | + | + | + |
| chr2R | 2421001 | 2500000 | + | + | + |  |
| chr2R | 2508001 | 2546000 | + | + | + |  |
| chr2R | 2551001 | 3506000 | + | + |  | + |
| chr2R | 3513001 | 3871000 | + | + | + | + |
| chr2R | 4414001 | 4425000 | + | + |  |  |
| chr2R | 4871001 | 5057000 | + |  |  |  |
| chr2R | 6283001 | 6476000 | + | + |  |  |
| chr2R | 10623001 | 10637000 | + |  |  | + |
| chr2R | 23133001 | 23313000 | + |  |  |  |
| chr3L | 4850001 | 5034000 | + |  |  |  |
| chr3L | 5043001 | 5075000 | + |  |  |  |
| chr3L | 13559001 | 13805000 | + |  |  |  |
| chr3L | 15196001 | 15475000 | + |  |  |  |
| chr3L | 18190001 | 18431000 | + |  |  |  |
| chr3L | 22577001 | 22627000 | + |  |  |  |
| chr3L | 22803001 | 22820000 | + |  |  |  |
| chr3L | 23157001 | 23173000 | + |  | + |  |
| chr3L | 23355001 | 23538000 | + |  |  |  |
| chr3L | 23550001 | 23679000 | + |  |  |  |
| chr3L | 23775001 | 24005000 | + |  |  |  |
| chr3L | 24056001 | 25075000 | + | + | + |  |
| chr3L | 25135001 | 25838000 | + | + | + |  |
| chr3L | 25844001 | 25965000 | + |  | + |  |
| chr3L | 26085001 | 26161000 | + |  | + |  |
| chr3L | 26166001 | 26311000 | + | + |  |  |
| chr3L | 26315001 | 26705000 | + | + | + |  |
| chr3L | 27391001 | 27815000 | + |  |  | + |
| chr3L | 28042001 | 28110227 | + |  |  |  |
| chr3R | 1 | 1257000 | + | + | + | + |
| chr3R | 1266001 | 1655000 | + | + |  |  |
| chr3R | 1664001 | 2569000 | + | + | + |  |
| chr3R | 2572001 | 2824000 | + | + | + | + |
| chr3R | 2831001 | 3034000 | + | + | + | + |
| chr3R | 3040001 | 3129000 | + | + |  | + |
| chr3R | 3136001 | 3533000 | + | + | + |  |
| chr3R | 3674001 | 3692000 | + | + |  |  |
| chr3R | 3827001 | 3842000 | + | + |  |  |
| chr3R | 3890001 | 4175000 | + |  | + |  |

**Supplementary Table 1 cont.**
|  |  |  |  |  |  |  |
| --- | --- | --- | --- | --- | --- | --- |
| chr3R | 6159001 | 6327000 | + |  |  |  |
| chr3R | 6515001 | 6624000 | + |  |  |  |
| chr3R | 7572001 | 7714000 | + |  | + |  |
| chr3R | 10919001 | 11142000 | + |  | + |  |
| chr3R | 16712001 | 16948000 | + |  |  |  |
| chr3R | 19704001 | 19715000 | + |  |  |  |
| chr3R | 22142001 | 22303000 | + |  |  | + |
| chr3R | 28020001 | 28252000 | + |  |  |  |

**Supplemental Table 2.**

half max position
| *Wild type* |  | max (log2) | left arm (bp) | right arm (bp) | half max total (bp) | fold change relative to wild type |
| --- | --- | --- | --- | --- | --- | --- |
| 7F | *chrX* | 3.911 | 8439400 | 8518500 | 79100 | 1 |
| 22B* | *chr2L* | 1.961 | 1888300 | 1937200 | 48900 | 1 |
| 30B | *chr2L* | 1.798 | 9504800 | 9587600 | 82800 | 1 |
| 34B | *chr2L* | 2.33 | 13371800 | 13438500 | 66700 | 1 |
| 62D | *chr3L* | 1.678 | 2231600 | 2314900 | 83300 | 1 |
| 66D | *chr3L* | 5.494 | 8694700 | 8765800 | 71100 | 1 |

half max position
| *SuUR-* |  | max (log2) | left arm (bp) | right arm (bp) | half max total (bp) |  |
| --- | --- | --- | --- | --- | --- | --- |
| 7F | *chrX* | 3.528 | 8425000 | 8528900 | 103900 | 1.313527181 |
| 22B* | *chr2L* | NA | NA | NA | NA | NA |
| 30B | *chr2L* | 1.682 | 9506100 | 9607700 | 101600 | 1.22705314 |
| 34B | *chr2L* | 2.355 | 13364800 | 13473500 | 108700 | 1.629685157 |
| 62D | *chr3L* | 1.745 | 2221200 | 2342400 | 121200 | 1.454981993 |
| 66D | *chr3L* | 4.88 | 8685700 | 8778600 | 92900 | 1.306610408 |

half max position
| *Rif1-* |  | max (log2) | left arm (bp) | right arm (bp) | half max total (bp) |  |
| --- | --- | --- | --- | --- | --- | --- |
| 7F | *chrX* | 3.824 | 8420700 | 8540700 | 120000 | 1.517067004 |
| 22B* | *chr2L* | NA | NA | NA | NA | NA |
| 30B | *chr2L* | 1.807 | 9496800 | 9607300 | 110500 | 1.334541063 |
| 34B | *chr2L* | 2.474 | 13361400 | 13474400 | 113000 | 1.694152924 |
| 62D | *chr3L* | 1.719 | 2217900 | 2335500 | 117600 | 1.411764706 |
| 66D | *chr3L* | 5.465 | 8679700 | 8783500 | 103800 | 1.459915612 |

\*22B is a strain specific amplicon present in Oregon R and not *SuUR* and *Rif1* mutants

